# Emergent Spatiotemporal Population Dynamics with Cell-Length Control of Synthetic Microbial Consortia

**DOI:** 10.1101/2021.04.06.438650

**Authors:** James J. Winkle, Bhargav R. Karamched, Matthew R. Bennett, William Ott, Krešimir Josić

## Abstract

Increased complexity of engineered microbial biocircuits highlights the need for distributed cell functionality due to concomitant increases of metabolic and regulatory burdens imposed on single-strain topologies. Distributed systems, however, introduce additional challenges since consortium composition and spatiotemporal dynamics of constituent strains must be robustly controlled to achieve desired circuit behaviors. Here, we address these challenges with a modeling-based investigation of emergent spatiotemporal population dynamics that result from cell-length control of monolayer, two-strain bacterial consortia. We demonstrate that with dynamic control of a strain’s division length, nematic cell alignment in close-packed monolayers can be destabilized. We found this destabilization conferred an emergent, competitive advantage on smaller-length strains—but by mechanisms that differed depending on the spatial patterns of the population. We used complementary modeling approaches to elucidate underlying mechanisms: an agent-based model to simulate detailed mechanical and signaling interactions between the competing strains and a reductive, stochastic lattice model to represent cell-cell interactions with a single rotational parameter. Our modeling suggests that spatial strain-fraction oscillations can be generated when cell-length control is coupled to quorum-sensing signaling in negative feedback topologies. Our research employs novel methods of population control and points the way to programming strain fraction dynamics in consortial synthetic biology.

Engineered microbial collectives are more versatile and robust than single strain populations. However, the function of such collectives is sensitive to their spatiotemporal organization. Here, we demonstrate control of the spatiotemporal composition of synthetic microbial consortia by dynamically modulating the average cell length of constituent strains. Such modulation confers an emergent “mechanical fitness” advantage upon the shorter length strain. We used both a biophysically realistic agent-based model to test the impact of cell shape on spatiotemporal dynamics and a conceptually simpler stochastic lattice model to explain the essential mechanisms driving the dynamics.

## Introduction

Understanding and designing microbial consortia with distributed functionality is of increasing interest in synthetic biology [6,20,28,31,43,50,53]. Assigning different functions to separate strains in a consortium reduces the metabolic load on each strain and thus allows more complex functionality and greater robustness [2,8,10,51,54]. Synthetic genetic circuits previously engineered in single strains, such as feedback oscillators and toggle switches [17,22], have recently been implemented in consortia [2,10,18,33]. However, we still lack the mathematical and computational tools that allow us to help engineer such systems in a principled way.

Synthetic microbial consortia intrinsically require balance and control of population strain fractions to acheive desired genetic circuit functionality. Important population control studies of synthetic bacterial collectives in the literature have employed both theoretical and experimental approaches, and a number of different control mechanisms have been introduced. Such approaches include predator-prey systems [5], cross-feeding auxotrophs [32], toxin-antitoxin [49] and “ortholysis” [53] mechanisms, and external switching control [52]. Although many of these methods employ regulating feedback, population dynamics achieved via toxic agents released by a cell must itself be tightly controlled to prevent unwanted expression and non-robust population behaviors. Studies that focus on distributed microbial systems have also been wide-ranging and include those of information exchange between constituent strains [26], ecological dynamics [12,46,63], metabolic resource allocation [27,44], microbial social interactions [37], and the human microbiome [39,55]. In each of these examples, balance and control of constituent parts is central to robust functionality [38,58].

In contrast to population control mechanisms employing a toxin, here we suggest that microbial consortium’s strain distribution can be controlled by changing the average division length of cells within each strain. Our approach stems from two active research areas in bacterial synthetic biology. One area focuses on how cell aspect ratio (cell length divided by cell width) affects cell ordering in close-packed environments [11, 14, 30, 42, 59, 62]. These studies have combined experimental, theoretical, and computational approaches to demonstrate that decreasing cell length generally decreases cell nematic ordering in spatially confined environments such as monolayer microfluidic devices. The second research area concerns *programming* bacterial cell aspect ratio by modulating expression of the cell-division proteins MreB and FtsZ [29,56,64]. Our modeling approach explores a synthesis of these two lines of research by proposing that cell division length can be modulated dynamically in a single experiment.

We simulated the growth, mechanical interactions and intercellular signaling of two microbial strains with different average cell lengths in a spatially-extended, monolayer microfluidic device. We considered populations of rod-shaped bacteria, whose axial growth leads to emergent columnar population structures [3, 30] and found that decreasing a cell’s average length can alter population dynamics by destabilizing this emergent columnar organization. Using an agent-based modeling (ABM) approach, we found that this mechanism gave shorter cells a competitive advantage (“mechanical fitness”) in close-packed microfluidic trap simulations. To better understand the essential dynamical mechanisms behind the emergent dynamics, we also developed a complementary lattice model (LM) approach based on the key features of the microbial growth and interactions [30]. For our LM, we mapped the cell division length to a single parameter—the probability of cell rotation upon division—which we hypothesized controlled the emergent population patterns seen in our simulations

## Results

### Agent-based modeling

#### Single-strain nematic cell ordering

To determine how cell length influences the spatiotemporal dynamics of bacterial populations, we used an agent-based model (ABM) of *E.coli* cells growing and dividing in an open-walled microfluidic device (see Methods). We began with single-strain simulations to measure steady-state cell ordering in two regions of the trap (bulk and edge) as shown in Fig 1. Each simulation was initiated by seeding the trap with 32 cells, randomly placed within the trapping region [3]. Following a trap-filling transient, cells in the bulk region formed nematically aligned vertical columns, as previously reported in modeling and experimental studies [7,14,30,33,59,60]. The ordering of cells in each region depended, however, on the average division length, 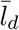, of the cell: Ordering in both regions decreased with decreasing 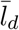 (Fig 1B) but maximal ordering in the bulk region required a minimum 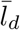. We define the order parameter as

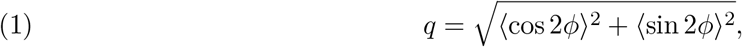

where *ϕ* denotes cell angle from horizontal in the lab frame and 〈·〉 signifies a region (bulk or edge) average. Correlation between cell length and nematic ordering has been reported previously [11,21,42].

**Figure 1.**
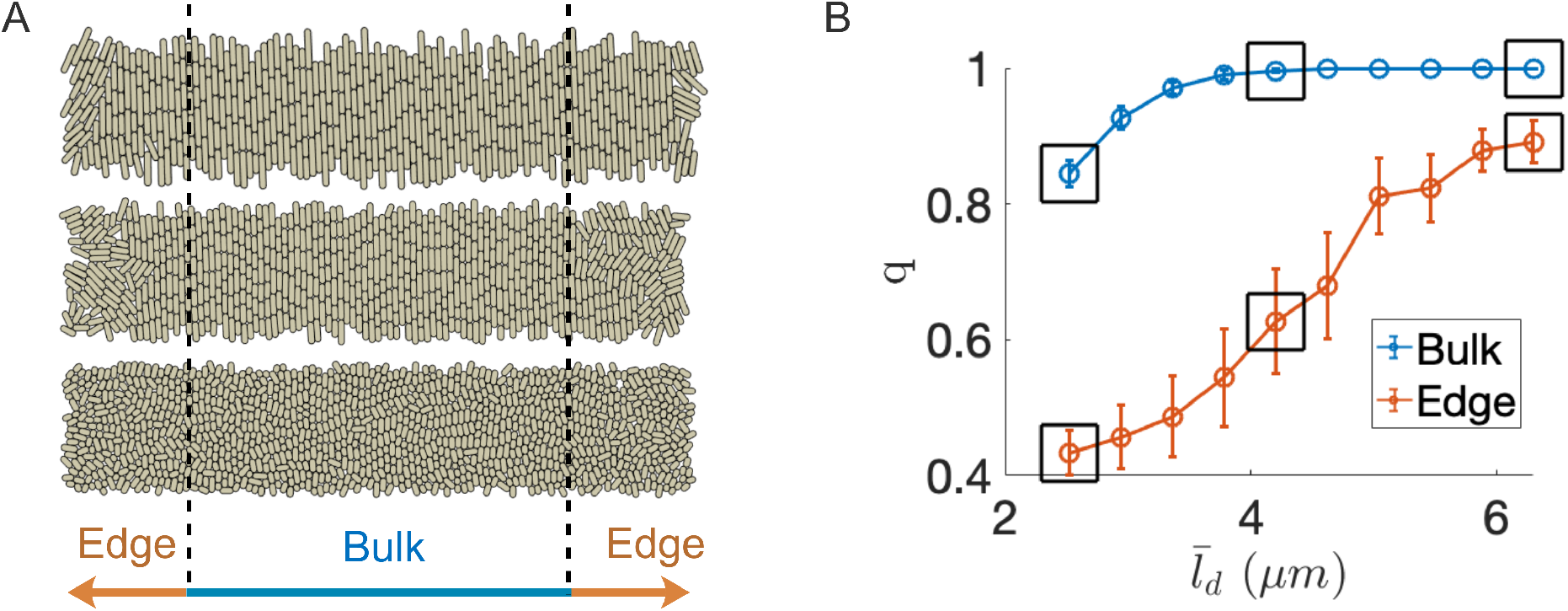
Average cell length affects nematic ordering in open-walled microfluidic trap simulations. (A) ABM simulation snapshots for three different average cell division lengths (2.5, 4.2, and 6.3*μ*m, bottom to top) in a 20 x 100 *μ*m open-walled microfluidic trap. For each simulation snapshot the order parameter, *q*, was measured in two different trap regions once the population reached steady-state (Eq. (1)). (B) Order parameter in the bulk and edge subregions. Circles and error bars represent means and standard deviations, respectively, for a single, representative simulation at each division length. Data was sampled twice per generation for 120 total samples after population stabilization (≈ 5 hours after cell seeding). With decreased average division length, cell ordering decreased in both regions, but disorder persisted in the edge regions even for the longest division lengths simulated. Boxed data corresponded to the three cases shown in panel (a).

For the smallest average division length we simulated (2.5 *μ*m, Fig 1A, bottom panel), the bulk population did not exhibit complete disorder (i.e., *q* > 0, where *q* = 0 would represent a uniform distribution of cell orientations [21]). However, we still observed cells that were oriented *horizontally* at random times and locations in each simulation run. We also observed horizontal cells in the bulk region for larger values of 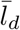, but not when 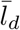 was greater than approximately 4 *μ*m. Horizontally oriented cells are significant since they can invade an adjoining column by axial growth. Based on these results, we hypothesized that horizontally oriented cells in the bulk region of a microfluidic population can alter population dynamics in two-strain consortia if one strain’s average division length is sufficiently small 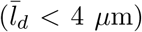 compared to the other 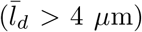, assuming a constant cell width of 1 *μ*m.

In multi-strain computational and experimental studies of morphologically homogeneous populations, columnar cell structure (nematic order) leads to long-term stabilization of the population ratios of each strain [3,60]. We thus conjectured that emergent nematic *disorder* could result in *destabilization* of columnar structure and thereby affect population ratio stability. To test this hypothesis, we performed ABM simulations of two-strain bacterial consortia where one strain’s average division length was reduced after the population structure stabilized with nematic order.

#### Well-mixed population bands and column invasion

To test the hypothesis that average cell length affects emergent columnar structure and therefore population dynamics in the close-packed environment of a microfluidic trap, we chose a ‘wild-type’ (WT) average division length of 4.2 *μ*m, a value for which (in single-strain simulations, see Fig 1) we observed nearly complete nematic ordering (*q* ≈ 1) in the bulk of the population in steady state. We then performed simulations with two strains and with two initial conditions: we used the WT strain and a ‘mutant’ strain (whose average cell length was reduced after population stabilization) in both random (Fig 2) and strain-separated (Fig 3) initial conditons.

**Figure 2.**
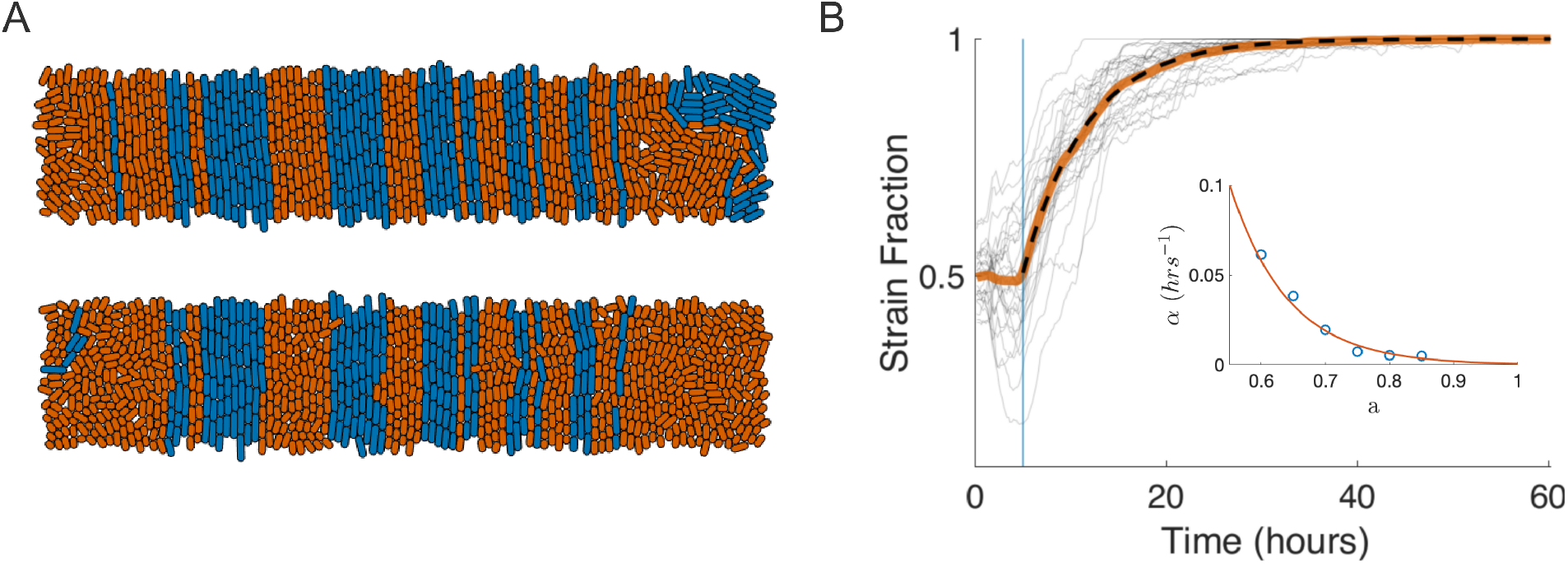
Cell morphology drives consortial population dynamics and strain fixation. (A) Two-strain ABM simulation snaphots in a 20 x 100 *μ*m trap. Top panel: With equal average cell division lengths 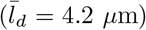, the consortium exhibited emergent columnar structure and stable strain population fractions in the bulk, as in Fig 1. Bottom panel: A snapshot taken approximately 5 generations (1.5 hours) after the orange strain’s average division length was reduced by a factor of *a* = 0.6 (resulting 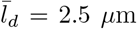). The increased rotational propensity of the smaller-length orange strain led to ejection of the blue strain by lateral invasion of its columns and subsequent growth-induced cell flow toward the open boundaries. Visible in lower panel are horizontal orange-strain cells beginning to destabilize adjacent blue-strain columns. (B) Orange strain fraction increased over time due to columnar invasion of the blue strain. At *t*≥ 5 hours (blue line), the orange strain’s average division length was reduced by factor *a* = 0.6. Destabilization of the columnar structure of the blue strain led to its eventual extinction in all simulations; Gray curves: 20 individual simulations; solid orange curve: mean strain fraction trajectory; dashed curve: Fit of 1 — 0.5*e*^-*α*(*t*-5)^. Inset: The fit rate parameter *α* decreases with length-reduction factor *a*.

**Figure 3.**
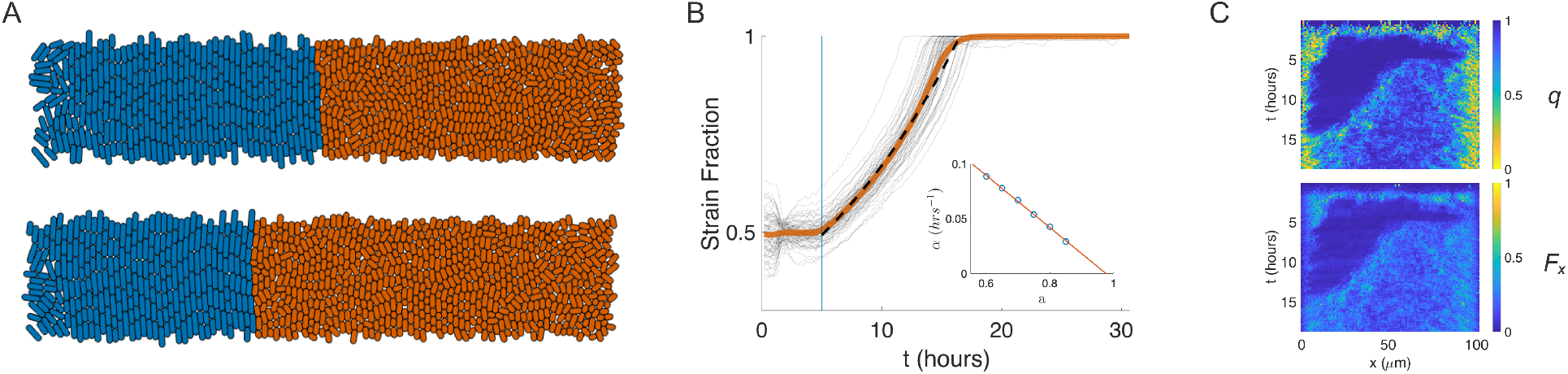
Single-interface consortial population dynamics and bulk forcing. (A) Snapshots of two-strain ABM simulation with cell strains seeded in separate halves of the trap. As in Fig 2, we reduced the mean division length of the orange cells to 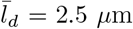 (*a* = 0.6) after stabilization of the population and emergence of nematic order in the bulk. Snapshots correspond to (top) one generation and (bottom) 10 generations (≈ 3 hours) after induction. (B) Orange strain population fraction time series for 20 ABM simulations. In contrast to the invasion mechanism illustrated in Fig 2, the orange strain here acted cooperatively by “bulk forcing”: the blue strain bulk population was pushed laterally and ejected out the left, open trap boundary. Gray curves: 20 individual simulations; solid orange curve: mean strain fraction trajectory; dashed curve: fit of 0.5*e*^*α*(*t*-5)^. Inset: the fit parameter *α* vs. the division length scale parameter, *a*. (C) Mechanisms of the bulk forcing are revealed by kymographs. Top panel: q order parameter averaged over 1*μ*m columns; Bottom panel: Growth-expansion force, *F_k_*, for each cell was projected horizontally to compute 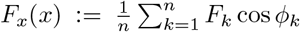, where *ϕ_k_* is the cell angle from horizontal and averaging is over cells *k* in a 1 *μ*m column *x*. Emergent disorder in the smaller length strain led to accelerating cumulative horizontal expansion force not present in the wild-type strain; the force imbalance ejected the longer length strain laterally.

In our first two-strain simulations, we seeded the trap with a total of 32, randomly placed, WT (blue) and mutant (orange) cells, where each strain type was selected with equal probability (see Fig 2B, WT strain colored blue). The two strains had identical growth rates [45] and were distinguished only by their average division lengths (see Methods). In the initial transient period of seed-cell growth and expansion (*t* < 5 hours), the two strains’ division lengths were identical. During this initial period, the strains’ population ratio stabilized after the formation of single-strain, columnar bands of various widths [3,60] (see Fig 2A, top image). With this emergent, nematic cell ordering, strain identity of each column was determined by the anchoring mother-cell position, which was located at or near the center of the column [3, 60].

At *t* = 5 hours (approximately 15 cell generations), we reduced the orange strain’s average division length by a factor of *a* = 0.6, which persisted through the remainder of the simulation. As a result of the division length decrease, nematic ordering markedly decreased for the shorter, orange strain (Fig 2A, bottom image), and as predicted by our single-strain simulations, orange cells randomly rotated into horizontal orientations. We then observed that horizontally oriented cells *invaded* the adjoining columns of the blue strain. When a cell rotated into an adjoining mother-cell position, subsequent growth and division of the invading cell allowed the invader and its descendants to occupy the invaded column by occupying the mother-cell position in the center of the column.

Fig. 2B shows the resulting temporal evolution of the orange strain fraction for 20 ABM simulations with the random cell-seeding condition for *a* = 0.6. We found that an exponential function of the form *f*(*t*) = 1 — 0.5*e*^-*α*(*t*-5)^ (dashed curve) provided an excellent description of the cell fraction (solid orange curve) average over these simulations. The observed exponential decrease in blue strain fraction has an intuitive explanation: If we view the loss of blue columns as a stochastic death process and assume that orange cells invade blue population bands at a constant rate *per blue column*, then the death rate is proportional to the number of remaining blue columns. We additionally found that the computed rate parameter for the exponential fit, *α*, itself depended exponentially on the average division length reduction factor, *a*. In the inset to Fig 2B, the exponential (orange curve) was fit to values of the rate paremeter *α* (blue circles) computed from similar ABM simulations per value of *a*. As expected from the data of Fig 1, *α* decreased as *a* increased due to decreased frequency of orange-strain cell rotations. However, why *α* depends exponentially on *a* remains to be explained.

#### Two-band populations and bulk rotational forcing

In the above two-strain ABM simulations we used a random initial seeding in space, which resulted in a banded population structure in steady state [3]. To understand the effect of the initial spatial patterning of strains on their subsequent population dynamics, we also seeded the trapping region so that the WT (blue) strain occupied the left half of the trap and the mutant (orange) strain the right half. This initializaton resulted in a single, initially stable strain interface near the center of the trap. However, in contrast to the previous ABM simulations, we did not observe frequent cell rotation and columnar invasion at the interface. Additionally, we observed that the orange strain’s population growth rate *increased* over time (Fig 3B, compare Fig 2B). We conjectured that a different mechanism was responsible for the population dynamics in the single-interface simulations than for those of the banded-interface simulations.

As in the previous simulations, after cell-length reduction was induced in the mutant (orange) strain, nematic disorder in that strain increased (as measured by the order parameter *q*, see Fig 3C). We observed with this initial condition, however, that the orange population did not frequently rotate horizontally at the single interface, but rather exerted a cooperative, horizontal force on the blue population due to the cumulative disorder exhibited in the *bulk* of orange strain. We then hypothesized that the observed increased rate of ejection of the WT strain was due to the increased probability that a mutant cell would grow and divide in a non-vertical direction (i.e, not in a vertically aligned state) once the bulk disorder in that strain was established and that the observed cooperative horizontal force would be proportional to the size of the bulk orange population. Hence, the force imbalance would lead to displacement and ejection of the blue strain, and the resulting rate of ejection would increase in time due to positive feedback.

To confirm our hypothesis that a second mechanism was responsible for the population dynamics observed in the separated initial condition, we measured both *q*, the order parameter, and *F_x_*, the column-averaged horizontal component of the cells’ growth-expansion force, and plotted them in kymographs. We define 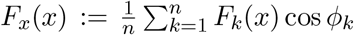, where *ϕ_k_* is the long-axis cell angle from horizontal in the lab frame, *F_k_* is the cell growth-expansion force [60], and we average over all cells *k* in a 1 *μ*m-wide column at horizontal location *x*. The decrease in *q*, following reduction of division length in the orange strain, is tightly correlated with an increase in *F_x_* in the bulk of the trap (Fig 3C,D). The resulting horizontal force imbalance between the two strains thus effected a net horizontal force on the bulk population of the blue strain. The magnitude of this force was indeed proportional to the size of the orange population, which resulted in positive feedback as originally hypothesized. We note that the observed persistent nematic disorder at the trap’s open boundaries are due to these boundaries being stress-free (see [60] and Fig 1).

We thus observed that population dynamics in single-interface simulations differed from those of the banded-interface in ways that confirmed the existence of a different strain-displacement mechanism. Figure 3B shows the evolution of the orange strain population-fraction again averaged over 20 ABM simulations (orange curve). An exponential, 0.5*e*^*α*(*t*-5)^, (dashed curve) provided a good fit to this average, up until the ejection of the blue strain. This exponentially increasing trend in the fraction of the WT train is consistent with the positive feedback mechanism we identified above. As we did for the banded simulations, we computed the rate parameter, *α*, for different values of the division-length scaling factor, *a* (Fig 3B, inset, circles). In contrast to the banded simulations, the fit rate parameter *α* depends approximately linearly on *a* in the single-interface case.

Thus, two different initial seeding conditions in the trap resulted in the displacement of the WT strain from the trap, but via two distinct mechanisms. However, it is not completely clear to what extent both mechanisms are present in both conditions: while it is easy to understand that bulk-forcing may cancel with a randomly distributed array of strain interfaces, we are not certain why we failed to observe columnar invasion in the single-interface simulations. One reason is that with multiple interfaces, there are simply more chances for an invasion to occur. Another potential reason is a decreased compliance of the WT strain with invasion by the blue strain: Emergent motion of the interface boundary may help the WT boundary column “escape” invasion events by transport at an advective timescale comparable to that of invasion events.

##### Lattice model

The ABM captures growth, cell-cell mechanical interactions, and diffusive signaling in realistic detail. As a result, however, it has many degrees of freedom, which make observed behaviors difficult to analyze in isolation and computationally expensive to simulate across even a narrow parameter space for each independent variable. Consequently, we used a complementary lattice model (LM) to understand the essential mechanisms by which cell morphology drives the spatiotemporal dynamics of consortia (See Methods). Since individual cells in the LM were simulated on a regular lattice, we used the rotation probability, *p*_rot_, as a proxy of cell size: As in the ABM, we assumed that shorter cells are less likely to change orientation, than their longer counterparts. ABM simulations showed that decreasing average cell division length confers fitness advantage via two distinct mechanisms, columnar invasion and lateral bulk forcing. Our simplified LM incorporates only the essential features of the ABM allowing us to test the hypothesis that the rotation probability alone facilitates invasion and forcing.

#### Impact of differing rotation probabilities in the LM

We hypothesized that assigning different rotation probabilities to the two strains in the LM will recapitulate the strain-strain interactions and the resulting spatiotemporal dynamics observed in the ABM. To verify this hypothesis, we considered two sets of LM simulations that mirrored those we performed with the ABM.

Before performing two-strain LM simulations, we verified that tuning the rotational probability *p*_rot_ in isogenic LM simulations reproduced the increase in nematic disorder with a reduction in average cell division length observed in the ABM. Figure 4A shows that in the LM the order parameter, *q*, computed as in the ABM, decreases, and cells are more likely to be oriented horizontally with an increases in *p*_rot_.

**Figure 4.**
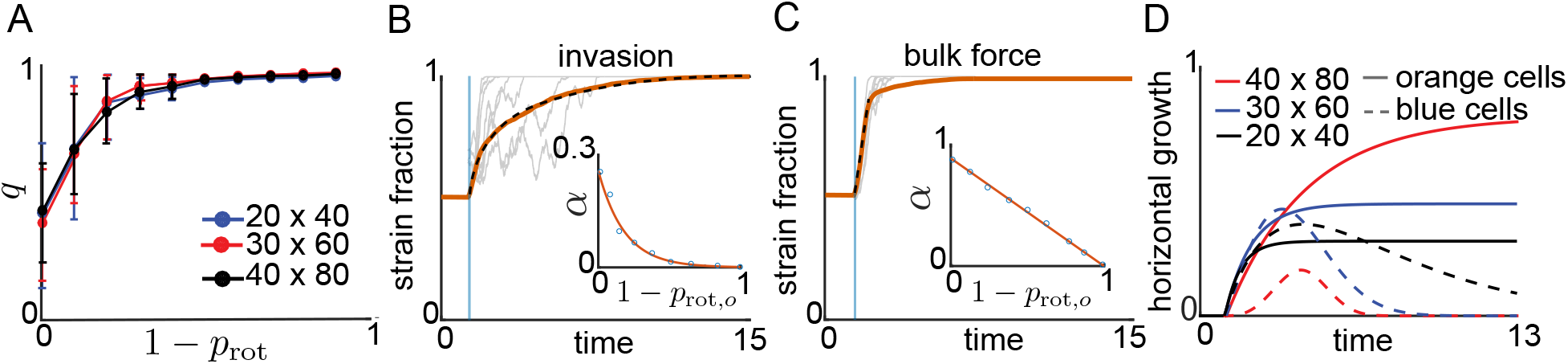
Lattice model simulations. (A) As in the ABM the order parameter, *q*, is obtained by averaging cell orientations across the trap at different times. Simulations with a single strain and different lattice sizes show that nematic disorder increases with *p*_rot_. (B) In two-strain LM simulations with multiple interfaces, we observed the same invasion mechanisms as in the ABM simulations. We initially set *p*_rot,*b*_(*t*) = *p*_rot,*o*_(*t*) = 0 so that during a transient period the two strains formed alternating stripes. Setting *p*_rot,*b*_(*t*) = 0 and *p*_rot,*o*_(*t*) = 0.5 at *t* 1 (light blue line), led to the ejection of the blue strain. The orange curve shows the evolution of the strain fraction averaged over 1000 trajectories, while gray curves show sample trajectories. The dashed curve shows the fit of the function 1 — 0.5*e*^-*α*(*t*-1)^ to the average orange strain fraction. Inset: The rate parameter, *α*, as a function of 1 — *p*_rot,*o*_ (circles represent LM data, compare to inset in Fig 2B.). (C) The LM also captured the bulk forcing mechanism we observed in the ABM. Simulations in (C) matched those in (B), with one change: We initially filled the left half of the lattice with orange cells, and the right half with blue cells. We fit the temporal evolution of the average orange strain fraction to 0.5*e*^*α*(*t*-1)^ over *t* ∈ [1, 2]. Inset: The rate parameter *α* depends linearly on 1 — *p*_rot,*o*_ (compare to inset in Fig 3B.) (D) With a single interface between the strains, horizontally growing cells forced out cells of the opposite strain, equivalent to the bulk forcing mechanism in the ABM. We plot the temporal evolution of the mean horizontal growth propensities for three lattice sizes with *p*_rot,*b*_ = 0.1 and *p*_rot,*o*_ = 0.5. Simulations in (C) corresponded to the red curves: The horizontal growth propensity of the orange strain dominates that of the blue strain, leading to ejection of the blue strain.

Our first set of two-strain LM simulations mirrored the multi-inference ABM simulations shown in Fig 2. We initialized each LM simulation by randomly assigning to each lattice site a cell of either the blue strain or the orange strain. Initial cell orientations were random (see Fig 7A). During the first phase of the simulation, we set *p*_rot,*b*_ (blue strain) and *p*_rot,*o*_ (orange strain) to zero to allow single-strain bands of columns of vertically oriented cells to emerge. At time *t* = 1, we set *p*_rot,*b*_ = 0 and *p*_rot,*o*_ = 0.5. Fig 4B shows the temporal evolution of the orange strain fraction, averaged over 1000 LM simulations (orange curve). The function *f* (*t*) = 1 — 0.5*e*^-*α*(*t*-1)^ again closely fits this average temporal evolution. When fit to data, the rate *α* varied approximately exponentially with 1 — *p*_rot,*o*_ (Fig 4B, inset). Importantly, the LM orange strain fraction dynamics and the dependence of *α* on 1 — *p*_rot,*o*_ closely matched the ABM orange strain fraction dynamics and the dependence of *α* on *a* (compare Fig 2B and 4B). This suggests that the LM successfully captures the invasion mechanism.

Our second set of two-strain LM simulations mirrored those of Fig 3. We initialized the lattice so that orange (blue) cells occupied the left (right) half of the trap, producing a single interface between the strains. To capture the decrease in division length upon induction, we set *p*_rot,*o*_ < *p*_rot,*b*_ at time *t* = 1. The increased rotational freedom in the orange strain allowed it to eject the blue strain through the right trap boundary, just as in the ABM simulations. In particular, the rate of increase of the orange strain fraction grew as the orange strain fraction increased over *t* ∈ [1,2], in accord with the positive feedback process observed in the ABM (see Fig 4C). We fit an exponential of the form 0.5*e*^*α*(*t*-1)^ to the mean orange strain fraction over *t* ∈ [1,2], and found that the rate parameter *α* changes approximately linearly with 1 — *p*_rot,*o*_ (Fig 4C, inset). This linear dependence was equivalent to that in the ABM simulations (compare Fig 3B and 4C) and suggests that the LM successfully captures the bulk forcing mechanism we observed in the ABM.

Unlike in the ABM, physical forces are not explicitly a part of the LM. However, upon division a horizontal or vertical cell moves a column of cells consistent with its orientation. Hence, the average horizontal growth probability describes the propensity of one strain to displace the other in the horizontal direction. We therefore defined the mean horizontal growth propensity for cells of strain *k* as

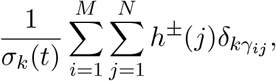

where *σ_k_*(*t*) is the number of cells of strain *k* in the lattice at time *t*, *h*^±^ is the horizontal growth rate, *δ_ij_* is the Kronecker delta function, and *γ_ij_* is the strain identity of the cell at the *i*^th^ row and *j*^th^ column in the lattice. Figure 4D shows the temporal evolution of the average horizontal growth propensities of each strain for the second set of LM simulations (red curves). The average horizontal growth propensity of the orange strain dominated that of the blue strain, explaining why the orange strain ejected its competitor.

We observed one important difference in the single-interface simulations between the LM and the ABM. In the ABM simulations, the movement of the interface between the strains accelerated until the blue strain was completely ejected from the trap. By contrast, a deceleration phase followed the initial acceleration phase in the LM simulations. This difference was due to a less stable interface between the strains in the LM compared to the ABM. In particular, we observed that stray blue cells remained in the trap even after the majority of the blue population was ejected. Nevertheless, the LM captured both mechanisms of WT displacement observed in the ABM remarkably well, despite being considerably simpler, and more tractable. Indeed, the LM can be described exactly by a master equation that can serve as a basis for further analysis.

##### Collective signaling: A spatial consortial oscillator

As multi-strain collectives grow and fill microfluidic devices, spatial structure emerges: Single-strain bands form of various widths, which depend on the cell seeding, but large-width bands can exceed the diffusion correlation length of interstrain communication [3]. We next suggest an experimentally feasible way to *control* such emergent spatial structures by using quorum-sensing (QS) signaling coupled to cell length modulation. In particular, we show computationally how oscillations in strain fraction can emerge when a QS signal from one strain induces a reduction in average division length in the opposite strain in a negative feedback, dual-state switch topology.

#### ABM strain fraction oscillator

We used our agent-based model (ABM) to show that spatiotemporal patterns in a microfluidic device can be controllably generated. To do so we combined a *bistable* QS circuit (based on the experimental circuit described in [2]) with the division-length reduction circuit we used in our two-strain simulations (see Fig 5). In this topology, each strain produces an orthogonal QS signal that activates the other strain’s LacI (a repressor protein) expression, which represses the production of QS signal in that strain. In our ABM simulations, when the QS signal received from the opposite strain surpassed a threshold concentration, *H_T_*, this also triggered the reduction in average cell division length, whiche se set to a fixed factor, *a* = 0.6. This is in contrast to our previous simulations where average division length was regulated by *exogenous* induction. In the oscillator ABM simulations, each QS signal was well-mixed across the trapping region and coupled via the flow channels (see Methods).

**Figure 5.**
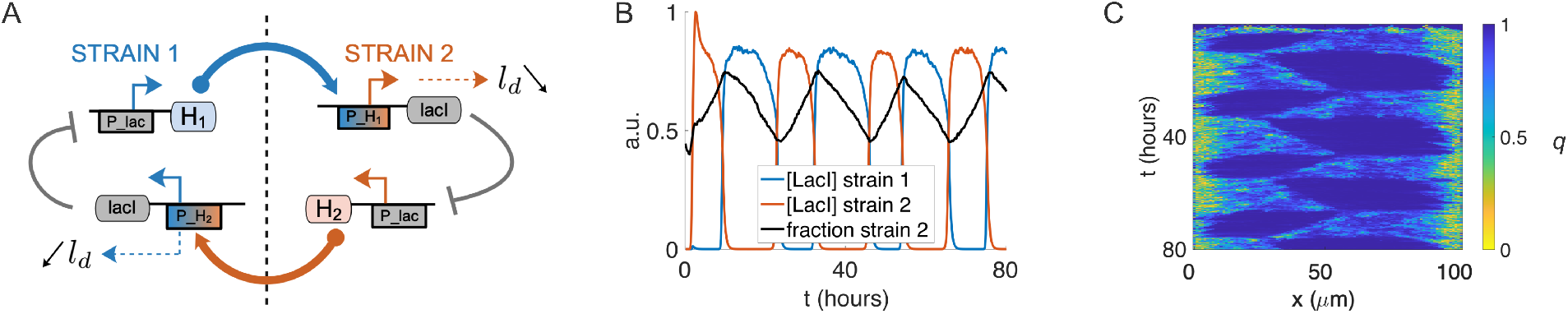
ABM oscillator. (A) Bi-stable circuit topology for quorum-sensing (QS) signal production, division length (*l_d_*) reduction, and QS repression by the protein LacI. (B) Sustained oscillations illustrated by the concentrations of LacI and strain fraction time series for an ABM simulation using the circuit topology in (a). Asymmetry in the strain fraction resulted from different QS molecules (C4HSL, C14HSL) with different diffusion rates. (C) Kymograph of order parameter *q* in the same simulation illustrating the switching behavior and emergent ordering dynamics (compare, Fig 3C).

As in Fig 3, we initialized these simulations by seeding the trap with 32 cells, with each strain occupying separate halves of the trap. Due to the random initial seeding, upon trap-filling each strain occupied approximately one-half of the trap, as in the simulations shown in Fig 3. We used published diffusion rates (see Methods) for the orthogonal QS molecules C4HSL and C14HSL [2], and assigned C14HSL production to the orange strain and C4HSL to the blue strain.

In the oscillator topology, we call the non-induced strain the “long” strain (i.e., it is the strain whose signaling is not repressed by LacI). The QS signal produced by the long strain reaches the other strain to repress its QS production and induce its division length reduction (see Fig 5A). As expected from the simulations shown in Fig 3, we observed in ABM oscillator simulations that the short strain cooperatively ejected the long strain via the bulk forcing mechanism. Thus, as the fraction of the long strain in the trap decreased, the QS signal received by the short strain decreased and eventually fell below *H_T_*, turning off both division-length reduction and its QS signal repression by negative-feedback. At this point QS production in the previously induced (short) strain switched on, triggering division length reduction and repression of QS signaling in the previously uninduced (long) strain. At this point the two strains exchange places, starting the next half-cycle in the oscillation.

This negative feedback topology produced sustained oscillations in the ABM. In Fig 5B, we plot the average LacI repressor concentration of each strain and the resulting strain fraction time series for an ABM simulation with division length reduction factor *a* = 0.6. A kymograph (Fig 5C) of the order parameter *q* (averaged as in Fig 3) shows the temporal transitions of cell ordering and resultant cooperative ejection of the majority strain in each oscillation half-period. We found that oscillations sustained indefinitely when the parameters were properly tuned, but with *H_T_* set too low, extinction of a strain would result (data not shown).

#### Reduced consortial oscillator model

To illuminate key features of the ABM consortial oscillator, we next describe a fast-slow dynamical system that captures the ABM consortial oscillator dynamics and their dependence on various timescales. In this effective model, the dynamics manifest as a relaxation oscillation. The driver of the oscillation is a state variable, *Q*, which represents the division-length state of the microbial strains. We let *Q* = 1 represent strain 1 (blue) being in a reduced division length state, while 1 — *Q* = 1 (i.e., *Q* = 0) represents the same for strain 2 (orange). Comparing with Fig 1A,B, *Q* is thus a proxy for the state of the QS promoter for LacI protein concentration, which is ON concomitantly with reduced division length. and determines the motion of the interface between the strains.

We simplified the trap geometry to one spatial dimension and model the strain-interface front position, *x* ∈ [0,1]. The dynamics for *Q* depend both on the proxy for concentration of strain 1 (respectively, strain 2) QS signals, which we assume proportional to *x* (1 — *x*), and to *Q* itself. In our reduced model we capture the QS sensitivity of each strain using ‘switching positions’: When *x* exceeds or falls below a specific value, denoted *K*_1,2_, the strains switch their division-length states through the action of an inverting Hill function, 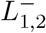, that represents the QS production promoter (repressed by LacI) in each strain. We thus capture the hysteresis of the toggle switch topology, Fig 5A, with the bistability of the switch state *Q* in the region *x* between switching locations.

Our reduced consortial oscillator model is

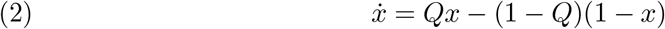

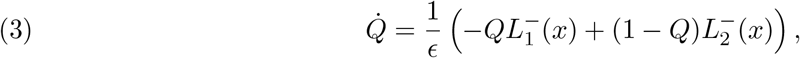

where 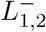 are decreasing Hill functions of 1 — *x* and *x*, respectively. The Hill functions are defined with QS switching locations at *x* = *K*_1_,*K*_2_ ∈ [0,1], such that

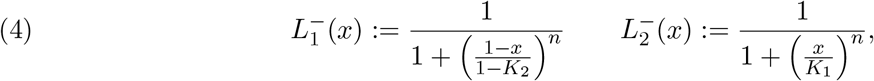

where *n* is the Hill exponent. The oscillations require *K*_1_ < *K*_2_, and *K*_2_ — *K*_1_ set to ensure separation of the timescales and sufficient switching time near the edges of the trap. Our reduced model assumes *x* dynamics are slow relative to *ϵ* (Eq. 3), which models the physiological timescale to effect changes of aspect ratio once the front position, *x*, approaches either of the values *K*_1_, *K*_2_. In the limit of large *n*, and small *ϵ*, the interface velocity, *x*, changes to positive (respectively, negative) as soon as *x* < *K*_1_ (*x* > *K*_2_). The values *K*_1,2_ thus define the two halves of the relaxation oscillation duty cycle. A simulation of Eqs. 2-3 shows good agreement with our ABM result (Fig 6C).

**Figure 6.**
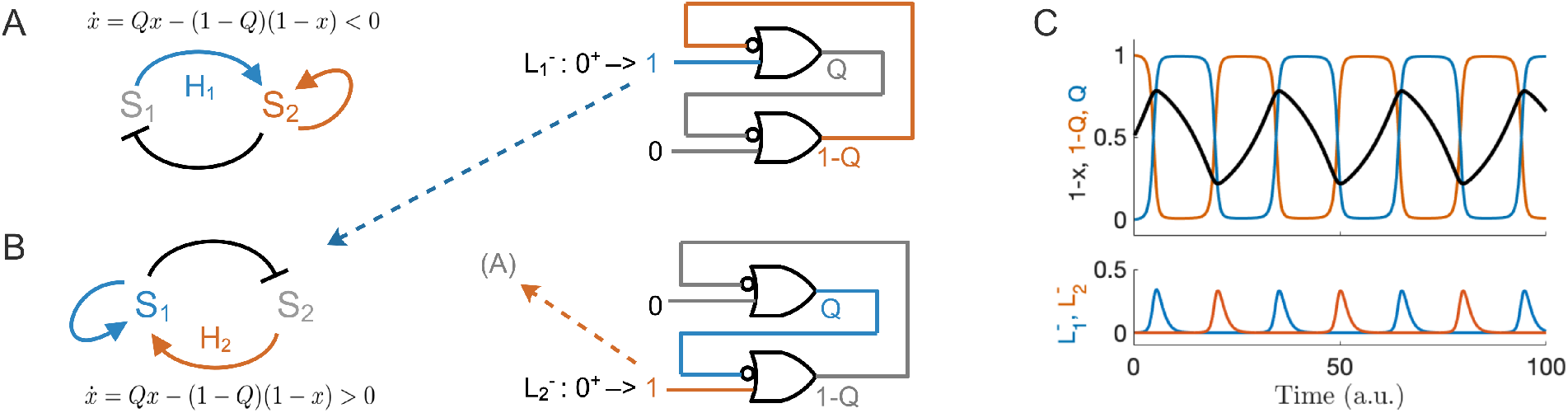
Consortial population relaxation oscillator. (A) Reduced oscillator model for *Q* = 0 (1 — *Q* = 1), which represents reduced cell length in strain 2. In this state, strain 2 represses strain 1 by negative motion of the strain interface *x* (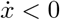 represents ejection of strain 1), which itself acts with positive feedback to increase the rate of motion. Signal *H*_1_ received from strain 1 activates the division length reduction and will diminish by negative feedback (since the strain 1 population is decreasing) eventually asserting the Hill function 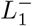 to change state to *Q* = 1. (B) Complementary model for state *Q* = 1. Reduced cell length in strain 1 represses strain 2 by positive motion of the interface *x* 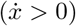 eventually asserting the Hill function 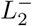 to revert the state to *Q* = 0, in turn. (C) Top panel: Interface position 1 — *x* and complementary states 1 — *Q* (orange) and *Q* (blue) plotted vs. time for *K*_1_ = 0.2, *K*_2_ = 0.8, and *n* = 8 (see Eqs. 2-3; cf. Fig 5B). Bottom panel: Hill functions 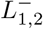 fire when the interface position reaches the respective thresholds, thereby switching the state of the memory circuit. The relaxation oscillator requires separation of the switching and front-travel timescales (see main text).

**Figure 7.**
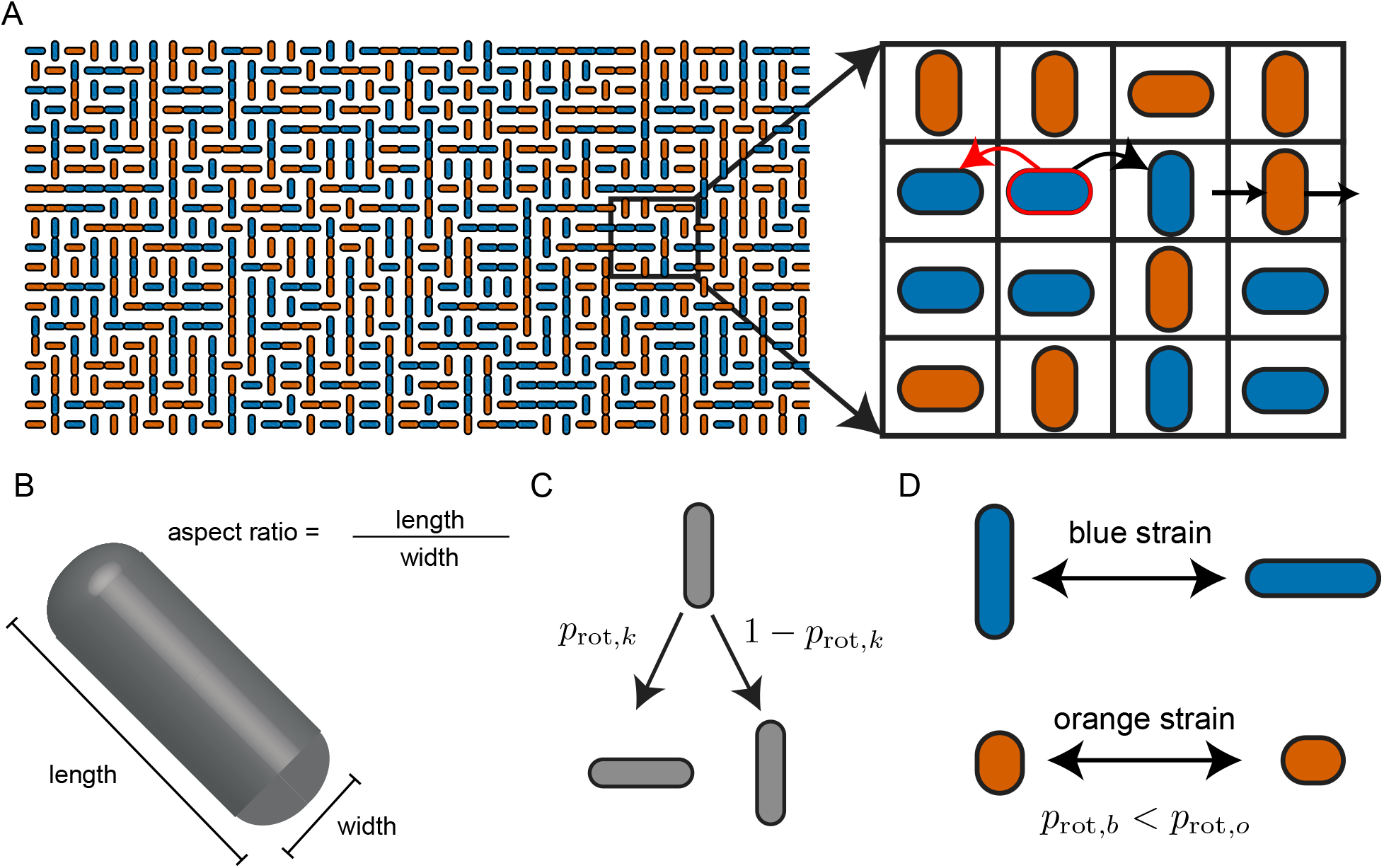
The lattice model (LM). **(A)** In the LM cell growth is directional and location-dependent: The horizontal cell outlined in red can grow to the right or left at location-dependent rates. The red arrow indicates that the leftward growth rate of the outlined cell is less than its rightward growth rate because this cell is located on the right side of the lattice. The black arrow indicates direction of cell division. When the cell divides, one daughter cell inherits the lattice position and orientation of the mother cell. The second daughter cell may rotate and occupies the lattice position immediately to the right of the mother cell, thereby displacing all existing cells in the direction of division by one unit. Cells that cross the lattice boundary are removed. **(B)** Schematic of a capsule-shaped bacterial cell. The cell’s aspect ratio is the ratio of its length to its width. **(C)** In two-strain LM simulations, the second daughter of a mother cell of strain type *k* rotates with probability *p*_rot,*k*_ if the 8 neighbors of the mother cell share her strain type. Otherwise, the second daughter rotates with a probability taken as the average of her probability of rotation and her 8 neighbors’ probabilities of rotation (see text for details). **(D)** In agreement with the effect of a smaller division length in the ABM, we assume that a smaller aspect ratio corresponds to higher rotation probability in the LM.

## Discussion

Controlling the spatiotemporal population dynamics of distributed, microbial systems is essential for optimizing their functionality. Here we demonstrated how bacterial cell morphology can be used to control spatiotemporal strain dynamics in the close-packed, monolayer environment of a microfluidic trap. We used both agent-based and lattice modeling to show that reducing average cell division-length in a two-strain consortium confers a “mechanical fitness” advantage to the shorter-length strain. This is in contrast to previous approaches to strain fraction control achieved primarily via exogenous signals [19,25], or induced cell lysis [40,61].

We showed a strain in a consortium can achieve a competitive advantage via two distinct mechanisms depending on the initial configuration of the strains. Using our agent-based model (ABM), we measured the order parameter, *q*, and observed its connection to the occurrence of horizontally oriented cells in the smaller length strain when two strains are intermingled in the trap, and increased lateral force imbalance when the two strains occupied different portions of the trap. Both configurations led to the ejection of the WT strain to the microfluidic trap open boundaries. The lattice model (LM) showed that both mechanisms can be explained by changes in a single parameter, the rotation probability, *p*_rot_. This parameter determined both the probability of invasion of a neighboring, opposite-strain column when the strains were intermingled, as well as the average lateral bulk force one strain exercised on the other, when the two strains occupied different parts of the trap.

Our ABM strain fraction oscillator and effective relaxation oscillation description showed that population control can be achieved using of a negative-feedback QS signaling topology in an experimentally relevant genetic circuit [2]. With a small modification to our circuit, stable strain fractions could be *programmed* by adding exogenous inducer to, for example, repress the division length protein expression in one of the strains. Similarly, by removing the division-length circuit altogether in one of the strains, a maximum population size could result from single-ended control in some range of initial conditions. We suggest that desired strain fractions could be robustly generated under a wide range of initial population distributions. Such control of consortium composition is a fundamental problem in synthetic biology [10,52], and earlier solutions relied primarily on the use of exogenous control [19, 25].

Understanding and controlling the behavior of distributed microbial systems is essential for engineering information exchange between constituent strains [26], ecological dynamics [12, 46, 63], metabolic resource allocation [27, 44], microbial social interactions [37]. The active control of cell morphology provides a way to engineer the spatiotemporal dynamics in synthetic systems, and to understand the patterns that emerge in microbial communities in nature.

## Methods

### Agent-based model

Our ABM captures mechanical interactions between cells growing in a microfluidic trap as well as the membrane and extracellular diffusion of signaling molecules produced intracellularly. We used this modeling framework to understand how changes in average cell length affect nematic cell ordering—and therefore emergent dynamics—in both homogeneous (single-strain) simulations and competing (two-strain) simulations. For two-strain simulations, we altered the mean division length of a strain with simulated external induction or by auto-regulatory control with quorum-sensing signaling topologies.

Cell growth, division, and induction. We modeled cell dynamics within a simulated, open-walled microfluidic device or *trap*. In ABM simulations, we assumed a monolayer rectangular trapping region with dimensions 20 × 100 *μ*m (height × width, see Fig 1–3). The trapping region is open on all sides to allow cell outflow. Two flow channels lie along the long edges of the device to provide nutrients and remove cells that exit the trapping region, thereby keeping the cell population approximately constant. Variants of this microfluidic design have been used in several published studies [3,10,13,33]. We also used quorum-sensing communication between strains and in this case modeled the flow channels as well-mixed compartments of signaling concentration. That is, we ignored directional effects of the flow for simplicity [10, 13, 33]. Our agent-based model thus simulated a population of growing and dividing *E. coli* cells in a 2D microfluidic trap environment using a mechanical interaction algorithm we described previously [60]. Here, we extended our previous approach to include bacterial quorum-sensing (QS) communication by integrating the finite-element software *Fenics* for numerical solution of the diffusion equation (see below).

Cells were modeled as 2D spherocylinders of constant, 1 *μ*m width. Each cell grew exponentially in length with a doubling time of 20 minutes [10, 33]. In order to prevent division synchronization across the population, when a mother cell of length *l* divided, the two daughter cells were assigned random birth lengths *ϵ*_0_*l* and (1 — *ϵ*_0_)*l*, where *ϵ*_0_ was sampled independently at each division from a uniform distribution on [0.45,0.55]. We set the division length of each daughter cell to 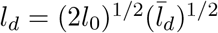, where *l*_0_ denotes the daughter’s birth length and 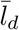 denotes the mean division length for the strain [4]. The mean division length, 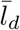, was modulated in a strain in our ABM simulations by simulated external induction or by using a quorum-sensing communication network between two strains. We divided the intracellular proteins and signaling molecules in the mother cell between the two daughters in proportion to their birth length.

For two-strain simulations, we initialized the ABM in two different ways and colored the strains orange and blue for identification (see Fig 2, 3) in simulation snapshots. We used in one case a spatially random initial seeding inside the trapping region (Fig 2). We chose the strain type of each seeded cell independently and with equal probability, which produced interchanging stripes of the two strains (Fig 2a) [3]. In another case we seeded the trapping region by placing blue cells in the left half and orange cells in the right half, which usually produced a single interface between the strains (Fig 3). In single-strain simulations, we used a neutral color for cell visualization (see Fig 1). Single-strain simulations were checked for strain fraction bias due to cell seeding with a negative control simulation (data not shown).

To investigate how cell length affects emergent dynamics in two-strain consortia, we reduced the mean division length (after a stabilization period for the population ratio) in the orange strain by a factor *a* ∈ [0.6,0.85], so that 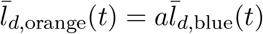 for *t* ≥ *t*_0_ (the width of the cells is the same for both strains for the duration of the simulations, see Fig 2, 3). Thus, the parameter *a* determined the difference in cell length between the two strains.

Quorum-sensing communication. To model inter-strain communication and feedback topologies for strain fraction control, we extended our ABM to include diffusive intercellular signaling by using the open-source finite element software *Fenics* [1,34,41]. We coupled intercellular signaling to the ABM by using an ODE for membrane diffusion of QS signals: For each cell of a strain, the intracellular QS concentration, *H*, depended on production at rate *α* (which in general depends on the signal coupling from the other strain), first-order membrane diffusion kinetics at rate *d*, and the local external QS concentration, *H_e_*. To model internal QS concentrations for a cell *i* we integrated the following differential equations for each QS molecule type in order to update both internal and external QS concentrations:

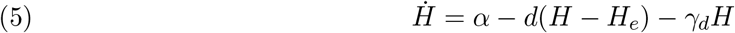

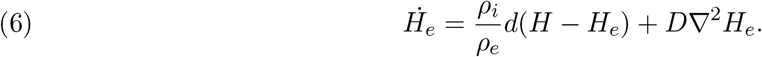

We thus assumed a well-mixed cell compartment and used the local extracellular concentration, *H_e_*(*x_i_, y_i_*), at the cell’s center, (*x_i_, y_i_*) in Eq (5). Here *γ_d_* is the cell dilution rate, *D* is the extracellular diffusion rate, and the dimensionless volume fraction ratio 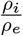 accounts for conservation of QS molecules in the local cell environment [10,33,47] (*ρ_i_* is the cell volume fraction, which we computed from simulations and set to 0.7, with *ρ_e_* = 1 — *ρ_i_*). All concentrations are generally functions of *x, y*, and *t*, which we omit to simplify notation.

We updated the external QS molecule concentrations over each time step by solving the 2D diffusion equation over the trapping region using the integrated *Fenics* solver. To simulate the perimeter of the trapping region and storage of signaling molecules in the flow channels, we used a splitting method whereby signal flux from the trapping region was integrated into a homogeneous channel volume, and diluted at rate *γ* due to media flow [10,13,33,48]. For two-strain consortia, we coupled signaling strength to cell-length control using a negative feedback topology. For strain 1, we reduced the mean division length of this strain when *H*_2_ > *H_T_*, where *H*_2_ is the measured intracellular QS signal from strain 2, and *H_T_* is a threshold value (similarly for Strain 2). Simulation code for our ABM and the integrated Fenics solver is open-source and available on Github.

### Lattice model

To illuminate the mechanisms that drive patterning in bacterial collectives, we developed a lattice model (LM) that captures the essential features of cell growth and strain-strain interactions. Lattice models have a storied history in biological modeling, and provide a valuable framework for modeling complex spatiotemporal dynamics in biological tissues [9,15,16,24,35,36, 57]. As is typical of lattice models, our LM gains tractability at the expense of some fidelity to reality.

We modeled the rectangular microfluidic trap as an *M*×*N* lattice in 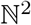. In the model, locations in the lattice were occupied by vertically or horizontally oriented cells belonging to one of two strains. Cells grew at location-dependent rates, as defined below. Upon division, one of the daughter cells replaced the mother cell, while the second daughter cell displaced a neighbor, and thus moved every cell in the direction of growth by one lattice site (see Fig 7a). We modeled traps with no walls, so any cell that crossed the boundary of the lattice disappeared from the system. Times between divisions in the trap were independent of one another, and exponentially distributed, with mean determined by the sum of growth rates of the cells in the trap. The locations of the division events were also independent of one another.

The assumption that cell division displaces the entire half-column or half-row in the direction of growth is strong. The impact of cell division may result in only local displacement [7, 42, 59, 60]. Nevertheless, we found that changing how many cells are moved after a cell division does not significantly alter qualitative behavior in our LM simulations.

We denote by *ν*^±^(*i*) the growth rate of a vertical cell in the *i*^th^ row toward the top (+) or bottom (–) boundary, and by *h*^±^(*j*) the growth rate of a horizontal cell in the *j*^th^ column toward the right (+) or left (–) boundary. We assumed that growth rates were modulated by the population that lies between a given cell and the closest boundary in the direction of growth, and hence that the growth rate of a vertical (horizontal) cell depended only on the row (column) in which it resided. We used a single parameter *κ* ∈ [0, ∞) to characterize how strongly the population modulated growth, and set

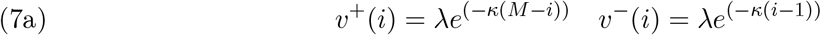

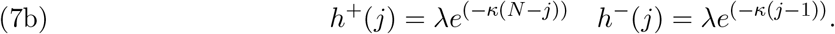

Using different decaying functions for *ν*^±^ and *h*^±^ did not alter LM dynamics significantly [30]. Since the growth rate determines the rate at which a cell divides we use the two terms interchangeably. Cell division and rotation. Previous simulations and experiments have shown that in crowded environments cell orientations evolve dynamically due to interactions with neighbors [7,11,30,42,59]. Rotational freedom increases as characteristic cell lengths decrease [11,56]. Our ABM simulations also showed that nematic disorder grows as mean cell division length decreases (See Fig 1). Since cells in the LM do not have physical dimensions, we introduced rotation probabilities, *p*_rot,*k*_, that determined the likelihood that cells in strain *k* change orientation from vertical to horizontal, or vice versa. Therefore, based on our ABM results, and previous experimental observations, we assumed that the probability of rotation increased monotonically with a decrease in mean cell division length. The precise dependence of *p*_rot,*k*_ on cell morphology is complicated. For simplicity we assumed that the intrinsic rotation probability of a cell is 1 — *p*_rot,*k*_ = *a*, where *a* is the normalized average division length of the smaller strain. As we discuss below, this probability can be affected by a cell’s neighbors.

In our simulations, a cell rotation only occurred at the moment of cell division. When a mother cell divided, the daughter cell that occupied the lattice site vacated by the mother did not rotate. The orientation of the daughter cell that displaced a neighbor differed from that of the mother cell with probability *p*_rot,*k*_ when the mother cell and her eight neighbors shared the same strain type (see Fig 7C). In crowded environments, the rotation dynamics of a given cell depends on the morphology of surrounding cells: If the surrounding cells are less likely to rotate, and hence better aligned, the given cell will be less likely to rotate as well. To account for this, in two-strain LM simulations, we set the displacing daughter’s rotation probability to the average rotation probability of itself and its eight neighbors. We implemented this model using a Gillespie algorithm [23] with the following events and corresponding rates: A a vertical (horizontal) cell of strain *k* at location (*i,j*) in the lattice displaced a neighbor at location (*i,j* ± 1) (respectively (*i* ± 1, *j*)) by producing a copy of equal orientation with probability *ν*^±^(*i*)(1 — *p*_rot,*k*_)Δ*t* (respectively *h*^±^(*j*)(1 — *p*_rot,*k*_)Δ*t*) and of differing orientation with probability *ν*^±^(*i*)*p*_rot,*k*_Δ*t* (respectively *h*^±^(*j*)*p*_rot,*k*_Δ*t*). Our LM results coincided remarkably well with our ABM results (see Results and Discussion). As with ABM simulations, we assigned a unique color to each strain for visualization.

## Author Contributions

JJW conceived of the idea. BRK, JJW, MRB, WO, and KJ designed the research. BRK, JJW, WO, and KJ developed the analysis. JJW performed simulations of the ABM. BRK performed simulations of the LM. BRK, JJW, WO, and KJ wrote the paper.

## Acknowledgements

This work has been partially supported by National Institutes of Health grant R01 GM117138 (MRB, WO, KJ), National Science Foundation grant DMS 1816315 (WO), National Science Foundation grant MCB 1936774 (MRB), National Science Foundation grant DMS 1662290 (MRB), National Science Foundation grant DMS 1662305 (KJ), National Science Foundation grant MCB 1936770 (KJ), and Welch Foundation grant C 1729 (MRB).

The authors acknowledge the use of the Sabine Cluster and the advanced support from the Research Computing Data Core at the University of Houston to carry out the research presented here.

